# The Masc–PSI complex directly induces male-type *doublesex* splicing in silkworms

**DOI:** 10.1101/2025.03.07.642132

**Authors:** Tatsunori Kaneda, Noriko Matsuda-Imai, Hidetaka Kosako, Keisuke Shoji, Masataka G. Suzuki, Yutaka Suzuki, Takashi Kiuchi, Susumu Katsuma

## Abstract

The WZ sex determination system is found in a diverse range of animals, including lepidopteran insects. In the silkworm *Bombyx mori*, a lepidopteran model insect, females carry the W chromosome with the feminizing gene *Feminizer* (*Fem*), which is the source of a female-specific PIWI-interacting RNA (piRNA). The *Fem* piRNA–PIWI complex cleaves an mRNA encoding the masculinizing protein Masculinizer (BmMasc), resulting in the production of the female-type splice variant of *Bombyx mori doublesex* (*Bmdsx*), which is the master genetic switch of *B. mori* sex differentiation. In contrast, in males, BmMasc induces the production of the male-type *Bmdsx* splice variant (*Bmdsx^M^*). However, the molecular mechanism through which BmMasc transduces the masculinizing signal to the male-specific *Bmdsx* splicing event remains unknown. In this study, we showed that BmMasc physically interacts with *Bombyx mori* P-element somatic inhibitor (BmPSI), that is a RNA binding protein required for *Bmdsx^M^* expression. *BmMasc* overexpression resulted in the production of *Bmdsx^M^* in *B. mori* ovary-derived BmN-4 cells, but this induction was severely inhibited in *BmPSI*-knocked down cells, indicating that BmPSI is essential for the masculinizing activity of BmMasc. We also identified that BmMasc-containing protein complex was associated with a region of *Bmdsx* pre-mRNA spanning introns 2 to 4, particularly around the junctions of intron 2 with exon 3 and intron 3 with exon 4. Exons 3 and 4 of *Bmdsx* are female-specific units and are skipped in *Bmdsx^M^*. Together with a previous report of the binding of BmPSI to the CE1 sequence located in exon 4 of *Bmdsx*, the current results strongly suggest that the BmMasc–BmPSI complex is required for male-specific exon skipping in *Bmdsx* pre-mRNA through its binding to the female-specific *Bmdsx* introns and exons.

## Introduction

Many animal species utilize sex determination systems which depend on the sex chromosome composition. Lepidopteran insects exhibit a female-heterogametic WZ sex determination system (Traut et al., 2007). In the silkworm *Bombyx mori*, females possess W and Z sex chromosomes, whereas males have two Z chromosomes. Genetic studies have demonstrated that the W chromosome alone determines femaleness in *B. mori* (Hasimoto, 1933), strongly suggesting that the dominant female-determining gene resides on the W chromosome. In 2014, we identified the primary female-determining gene, *Feminizer* (*Fem*), on the W chromosome of *B. mori* (Kiuchi et al., 2014). *Fem* encodes the precursor of a single PIWI-interacting RNA (piRNA), called *Fem* piRNA. The complex of *Fem* piRNA and *B. mori* PIWI protein SIWI cleaves the mRNA of the Z-linked masculinizing gene *B. mori Masculinizer* (*BmMasc*) (Kiuchi et al., 2014). Knockdown and knockout studies revealed that *Masc* homologs are essential for masculinization and dosage compensation in lepidopteran insects (Kiuchi et al., 2014; Lee et al., 2015; Fukui et al., 2015; Xu et al., 2017; Fukui et al., 2018; Wang et al., 2019; Harvey-Samuel et al., 2020; Deng et al., 2021; Visser et al., 2021; Bi et al., 2022; Tomihara et al., 2022; Pospíšilová et al., 2023; Li et al., 2024, Moronuki et al., 2024; Van’t Hof et al., 2024). In *B. mori*, for example, embryonic knockdown of *BmMasc* leads to the production of the female-specific isoform of *B. mori doublesex* (*Bmdsx^F^*) in male embryos (Kiuchi et al., 2014), indicating that BmMasc is essential for the expression of the male-specific isoform of *Bmdsx* (*Bmdsx^M^*).

The *Bmdsx* gene encodes a transcription factor that functions at the downstream end of the *B. mori* sex determination cascade. *Bmdsx* pre-mRNA undergoes alternative splicing in a sex-dependent manner (Ohbayashi et al., 2001), and the resulting BmDSX^F^ and BmDSX^M^ proteins are essential for sexual differentiation in *B. mori* (Suzuki et al., 2003; Suzuki et al., 2005). Previous studies that used cultured cells derived from *B. mori* male and female embryos identified two factors involved in sex-specific splicing of *Bmdsx* (Suzuki et al., 2008; Suzuki et al., 2010). BmPSI, a *B. mori* homolog of P-element somatic inhibitor, was shown to bind to the CE1 sequence in exon 4 of *Bmdsx*, which is excluded in *Bmdsx^M^*. Knockdown of *BmPSI* in male embryo-derived cultured cells resulted in increased *Bmdsx^F^* expression, suggesting the involvement of BmPSI in *Bmdsx^M^* production (Suzuki et al., 2008). Moreover, BmIMP, a *B. mori* homolog of IGF-II mRNA binding protein (IMP), was identified as the factor that enhances the BmPSI’s RNA binding activity to the CE1 sequence (Suzuki et al., 2010). However, it remains unknown how the primary male determiner BmMasc transmits its signal to the sex-specific splicing of *Bmdsx*.

Here, we performed BmMasc interactome analysis using *B. mori* ovary-derived BmN-4 cells and identified BmPSI as a candidate BmMasc-interacting protein. Subsequent experiments revealed that the BmMasc–BmPSI complex was required for BmMasc-induced expression of *Bmdsx^M^* through its binding to a region adjacent to the female-specific exons in *Bmdsx* pre-mRNA.

## Results

### Identification of BmPSI as the BmMasc-interacting protein

We previously reported that transient expression of GFP-fused BmMasc (BmMasc-GFP) or GFP-fused BmMascΔNLS (BmMascΔNLS-GFP, which is a BmMasc derivative lacking a nuclear localization signal) induces the expression of *Bmdsx^M^* in *B. mori* ovary-derived BmN-4 cells (Sugano et al., 2016). This strongly suggests that the factors interacting with both BmMasc-GFP and BmMascΔNLS-GFP are involved in the BmMasc-induced masculinizing pathway. To identify the components of the BmMasc-dependent masculinizing complex, we overexpressed BmMasc-GFP, BmMascΔNLS-GFP, or GFP (control) in BmN-4 cells, collected and lysed the transfected cells, and performed co-immunoprecipitation using anti-GFP nanobody-conjugated magnetic agarose beads. The proteins co-immunoprecipitated with BmMasc-GFP, BmMascΔNLS-GFP, or GFP, were subjected to LC-MS/MS analysis. Candidate proteins for interaction with both BmMasc-GFP and BmMascΔNLS-GFP were identified, which included various proteins potentially that are involved in RNA splicing (Fig. 1A, Table S1). Among them, we focused on BmPSI, a protein that was previously characterized as a regulator of *Bmdsx* male-type splicing (Suzuki et al., 2008), which interacted with both BmMasc-GFP and BmMascΔNLS-GFP (Fig. 1A). We further verified the interaction between exogenous BmMasc and endogenous BmPSI through immunoprecipitation of BmMasc-GFP-overexpressed cell lysate, followed by Western blotting with an anti-BmPSI antibody (Fig. 1B).

**Fig. 1.**
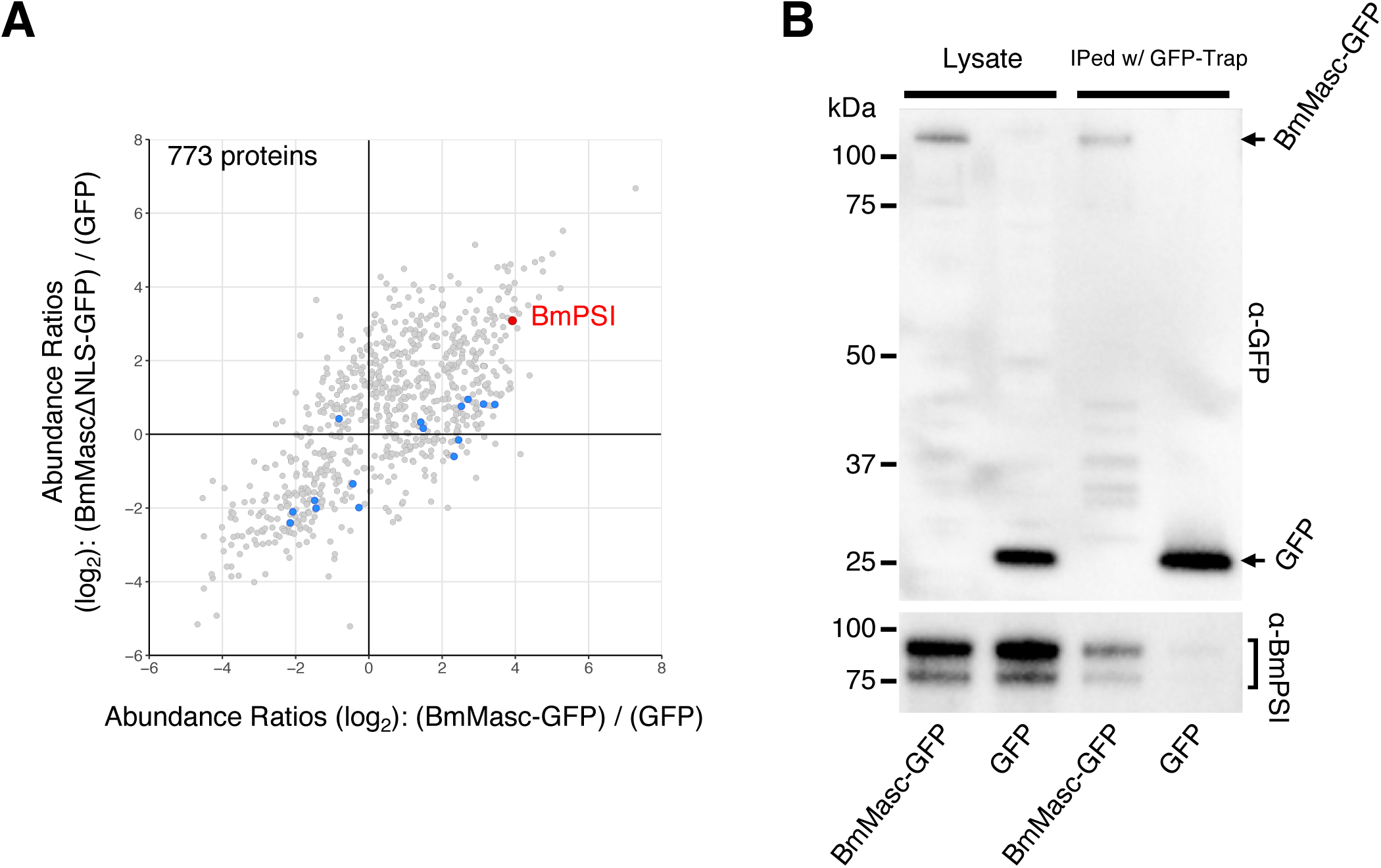
Identification of BmPSI as the BmMasc-interacting protein. (**A**) LC-MS/MS analysis of the immunoprecipitates obtained using anti-GFP nanobody-conjugated magnetic agarose beads from *BmMasc-GFP-*, *BmMascΔNLS-GFP-*, or *GFP-*transfected BmN-4 cells. The dots indicate the proteins detected in this analysis (773 proteins). BmPSI is indicated by red. The potential proteins involved in RNA splicing are shown in blue (see Table S1). The quantitative data are shown on a log-log scale. The x-axis represents the abundance ratio log_2_(BmMasc-GFP)/(GFP), and the y-axis represents the abundance ratio log_2_(BmMascΔNLS-GFP)/(GFP). (**B**) Co-immunoprecipitation experiments using BmN-4 cells transfected with *BmMasc-GFP* or *GFP*. The immunoprecipitates with anti-GFP nanobody agarose beads were immunoblotted using an anti-GFP or anti-BmPSI antibody. The BmPSI bands are indicated by a bracket. Similar results were obtained in two independent experiments.

Next, we investigated domains of BmMasc and BmPSI that mediate their interaction. First, two BmPSI-mCherry derivatives, N-BmPSI-mCherry and C-BmPSI-mCherry, each of which contained four KH domains (putative RNA-binding domains) and two AB motifs (likely involved in protein–protein interactions), respectively, were constructed (Fig. 2A). Western blotting of the immunoprecipitates revealed that C-BmPSI-mCherry interacted with BmMasc-GFP, whereas N-BmPSI-mCherry did not (Fig. 2B); this finding suggested that the BmMasc-interacting region of BmPSI localizes to its C-terminus. Further experiments using two additional AB motif mutants, dAB1-C-BmPSI-mCherry (without the first AB motif) and dAB2-C-BmPSI-mCherry (without the second AB motif) (Fig. 2A), showed that these mutants almost lost their ability to bind BmMasc-GFP (Fig. 2C), suggesting the importance of the two AB motifs. We also generated two BmMasc-GFP derivatives, dzf1-BmMasc-GFP and dzf2-BmMasc-GFP, which lacked either of the two CCCH zinc finger domains (Fig. 2D). Western blotting of the immunoprecipitates revealed that both derivatives bound to BmPSI (Fig. 2E), suggesting that the CCCH zinc finger domains of BmMasc are dispensable for the interaction with BmPSI. We further generated the BmMasc-GFP derivative, CS-BmMasc-GFP, which possessed amino acid substitutions in the masculinizing domain (Fig. 2D) (Katsuma et al., 2015) and lacked the masculinizing activity (Fig. 2F). We observed that this derivative possessed binding activity for BmPSI (Fig. 2G). These results indicate that known functional domains in BmMasc are not involved in its binding to BmPSI.

**Fig. 2.**
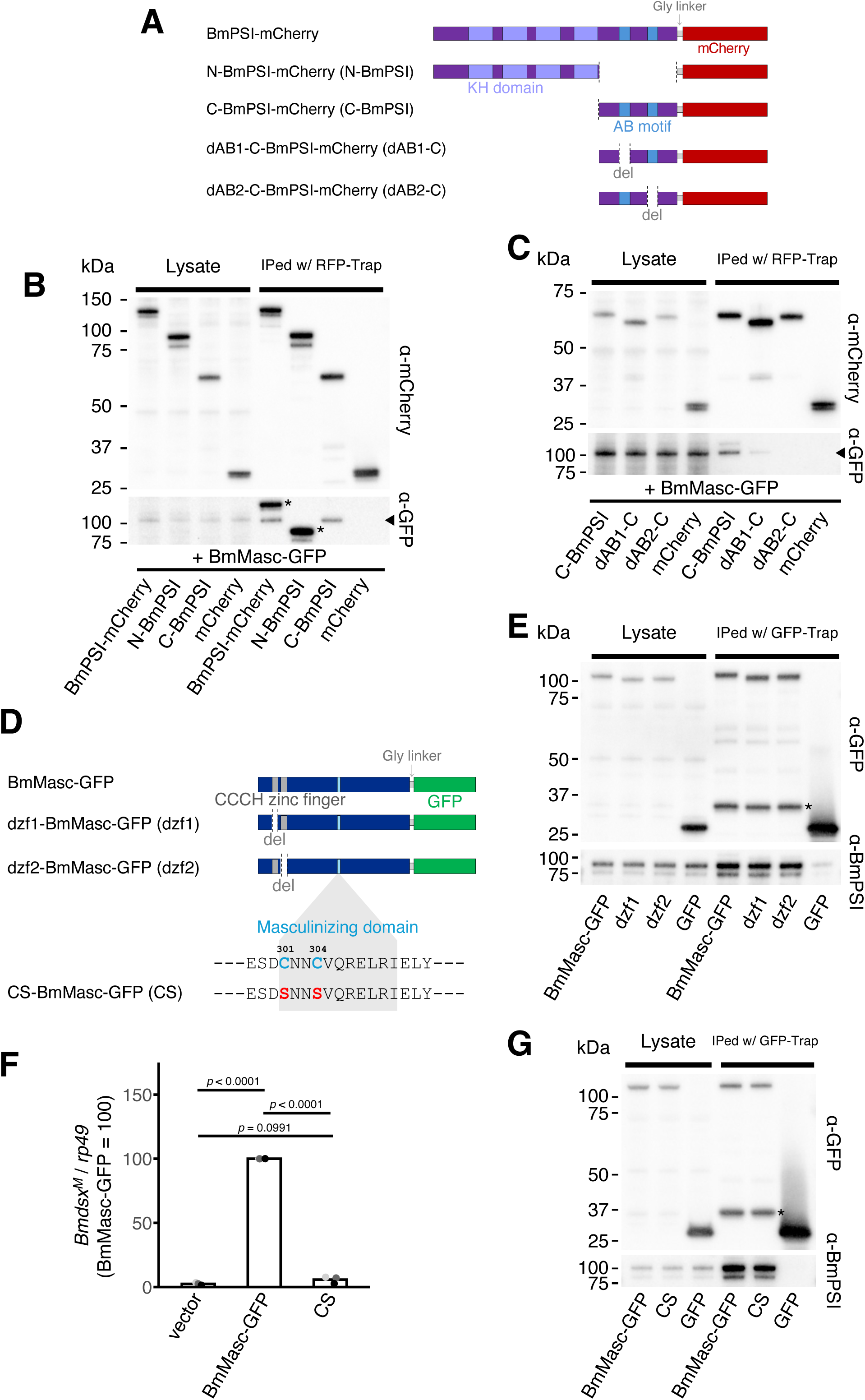
Identification of protein domains that mediate the interaction between BmMasc and BmPSI. (**A**) Structure of BmPSI-mCherry derivatives. The KH domains and AB motifs in BmPSI, glycine linker, and mCherry are shown. (**B**) Co-immunoprecipitation experiments using BmN-4 cells co-transfected with *BmMasc-GFP* and *BmPSI-mCherry*, *N-BmPSI-mCherry* (*N-BmPSI*), *C-BmPSI-mCherry* (*C-BmPSI*), or *mCherry*. The immunoprecipitates with anti-RFP nanobody agarose beads were immunoblotted using an anti-mCherry or anti-GFP antibody. The BmMasc-GFP bands are indicated by an arrowhead. Non-specific bands are indicated by asterisks. Similar results were obtained in two independent experiments. (**C**) Co-immunoprecipitation experiments using BmN-4 cells co-transfected with *BmMasc-GFP* and *C-BmPSI-mCherry* (C-BmPSI), *dAB1-C-BmPSI-mCherry* (dAB1-C), *dAB2-C-BmPSI-mCherry* (dAB2-C), or *mCherry*. The immunoprecipitates with anti-RFP nanobody agarose beads were immunoblotted using an anti-mCherry or anti-GFP antibody. The BmMasc-GFP bands are indicated by an arrowhead. Similar results were obtained in two independent experiments. (**D**) Structure of BmMasc-GFP derivatives. The CCCH zinc finger domains and masculinizing domain in BmMasc, glycine linker, and GFP are shown. In CS-BmMasc-GFP, the functionally important cysteine residues (Cys-301 and Cys-304) in the masculinizing domain are replaced by serine. (**E**) Co-immunoprecipitation experiments using BmN-4 cells transfected with *BmMasc-GFP*, *dzf1-BmMasc-GFP* (dzf1), *dzf2-BmMasc-GFP* (dzf2), or *GFP*. The immunoprecipitates with anti-GFP nanobody agarose beads were immunoblotted using an anti-GFP or anti-BmPSI antibody. Non-specific bands are indicated by an asterisk. Similar results were obtained in two independent experiments. (**F**) Expression of *Bmdsx^M^*in BmN-4 cells transfected with *BmMasc-GFP* or *CS-BmMasc-GFP* (CS). The *Bmdsx^M^* levels were estimated by RT-qPCR. The data are the means of three independent experiments. Adjusted *p* values from Tukey’s multiple comparisons tests are shown. (**G**) Co-immunoprecipitation experiments using BmN-4 cells transfected with *BmMasc-GFP* or *CS-BmMasc-GFP* (CS). The immunoprecipitates with anti-GFP nanobody agarose beads were immunoblotted using an anti-GFP or anti-BmPSI antibody. Non-specific bands are indicated by an asterisk. Similar results were obtained in two independent experiments.

### BmPSI is required for BmMasc-dependent *Bmdsx^M^* expression in BmN-4 cells

Unlike *BmMasc*, the exogenous overexpression of *BmPSI* alone did not induce *Bmdsx^M^* expression (Fig. 3A). This result is consistent with the finding that the levels of *BmPSI* mRNA are comparable between male and female cells (Suzuki et al., 2008). *BmMasc* is post-transcriptionally regulated by *Fem* piRNA in BmN-4 cells (Kiuchi et al., 2014) and was hardly detected at the protein level (Fig. 3B). Considering that BmPSI protein is abundantly expressed in BmN-4 cells (Fig. 3C), BmPSI likely functions as a component of the masculinizing complex by cooperating with the exogenously expressed BmMasc protein in BmMasc-transfected BmN-4 cells.

**Fig. 3.**
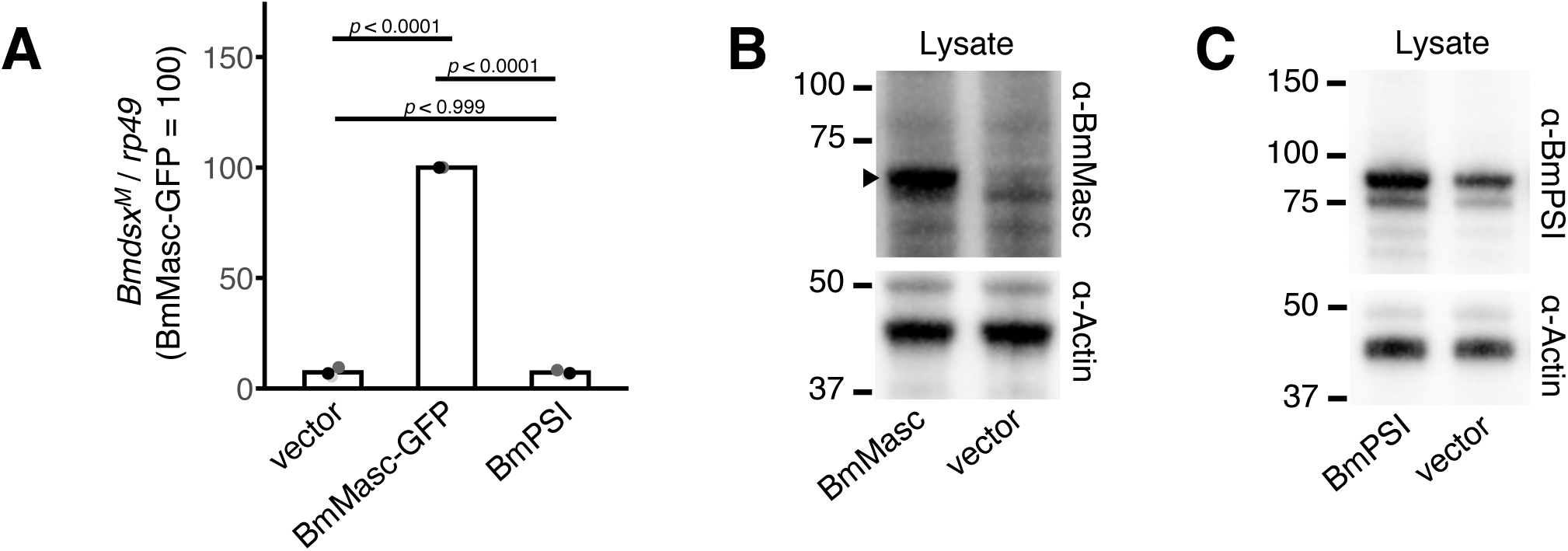
Abundance of BmPSI and BmMasc proteins in BmN-4 cells. (**A**) Expression of *Bmdsx^M^* in BmN-4 cells transfected with *BmMasc-GFP* or *BmPSI*. The *Bmdsx^M^* levels were estimated by RT-qPCR. The data represents the means of three independent experiments. Adjusted *p* values from Tukey’s multiple comparisons tests are shown. (**B**) Western blotting experiments of BmN-4 cell lysates transfected with *BmMasc* or empty vector. The cell lysates were immunoblotted using an anti-BmMasc or anti-actin (loading control) antibody. The BmMasc bands are indicated by an arrowhead. Similar results were obtained in two independent experiments. (**C**) Western blotting experiments of BmN-4 cell lysates transfected with *BmPSI* or empty vector. The cell lysates were immunoblotted using an anti-BmPSI or anti-actin (loading control) antibody. Similar results were obtained in two independent experiments.

To investigate whether BmPSI is required for BmMasc-dependent masculinization in BmN-4 cells, we knocked down *BmPSI* using double-stranded RNA (dsRNA) soaked in BmN-4 sid-1 cells. As shown in Fig. 4A and B, two types of dsRNAs targeting *BmPSI* (ds*BmPSI*-1 and ds*BmPSI*-2) both efficiently reduced the amount of BmPSI protein compared with the ds*Luciferase* (ds*Luc*)-treated (control) cells. We then transfected *BmMasc* cDNA into *BmPSI*-knocked down cells and assessed the level of BmMasc-dependent masculinization. The expression level of *Bmdsx^M^* was markedly lower in *BmPSI*-knocked down cells compared with the ds*Luc*-treated cells (Fig. 4C), indicating that BmPSI is essential for the BmMasc-dependent masculinizing pathway in BmN-4 cells.

**Fig. 4.**
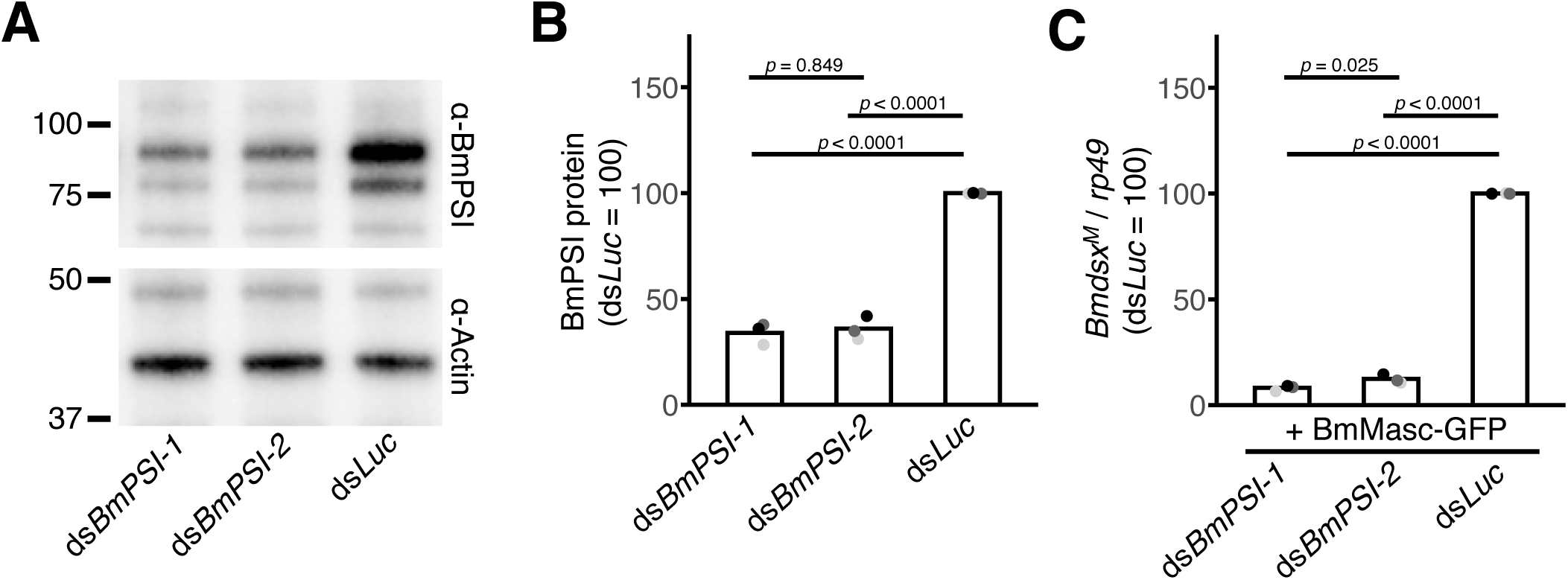
BmPSI is required for BmMasc-dependent *Bmdsx^M^* expression in BmN-4 cells. (**A**) Western blotting of dsRNA-treated BmN-4 sid-1 cell lysates. The cell lysates were immunoblotted using an anti-BmPSI or anti-actin (loading control) antibody. Similar results were obtained in three independent experiments. (**B**) Quantification of BmPSI protein abundance in BmN-4 sid-1 cells treated with ds*BmPSI-1*, ds*BmPSI-2*, or ds*Luc* (control). The BmPSI protein levels were estimated by Western blotting, as shown in (A). The data are the means of three independent experiments. Adjusted *p* values from Tukey’s multiple comparisons tests are shown. (**C**) Expression of *Bmdsx^M^* in BmN-4 sid-1 cells treated with ds*BmPSI-1*, ds*BmPSI-2,* or ds*Luc* (control). The *Bmdsx^M^* levels were estimated by RT-qPCR. The data are the means of three independent experiments. Adjusted *p* values from Tukey’s multiple comparisons tests are shown.

### The BmMasc–BmPSI complex physically interacts with *Bmdsx* pre-mRNA

It has been reported that BmPSI binds to the CE1 sequence in exon 4 of *Bmdsx* pre-mRNA and regulates *Bmdsx^M^* expression (Suzuki et al., 2008). We hypothesized that the BmMasc–BmPSI complex physically interacts with *Bmdsx* pre-mRNA to induce male-specific splicing. To test this hypothesis, we performed RNA immunoprecipitation (RIP) experiments using *BmMasc-GFP*-or *GFP*-transfected BmN-4 cells. Cell lysates were immunoprecipitated with anti-GFP beads, and RNA fragments were extracted from the immunoprecipitates and analyzed by RNA-seq and RT-qPCR (Fig. 5A). Mapping of the RNA-seq reads onto the genomic region of the *Bmdsx* gene revealed that RNA fragments associated with the BmMasc-GFP-containing protein complex were enriched in the regions spanning from the intron 2 to intron 4 of *Bmdsx* pre-mRNA, especially around the junction of intron 3 with exon 4 (Fig. 5B). RIP-qPCR experiments targeting the intron–exon junctions revealed the enrichment of RIP fragments around the junctions of intron 2 with exon 3 and intron 3 with exon 4 (Fig. 5C). RIP-qPCR amplicon for exon 4 contained the CE1 sequence, thus confirming the previous observation of BmPSI binding to the CE1 sequence (Suzuki et al., 2008). Based on these results, we conclude that the BmMasc–BmPSI protein complex serves as the core masculinizing machinery by binding to the regions located around the junctions of intron 2 with exon 3 and intron 3 with exon 4, inhibiting their splicing, and thereby promoting *Bmdsx^M^* expression in *B. mori* males (Fig. 5D).

**Fig. 5.**
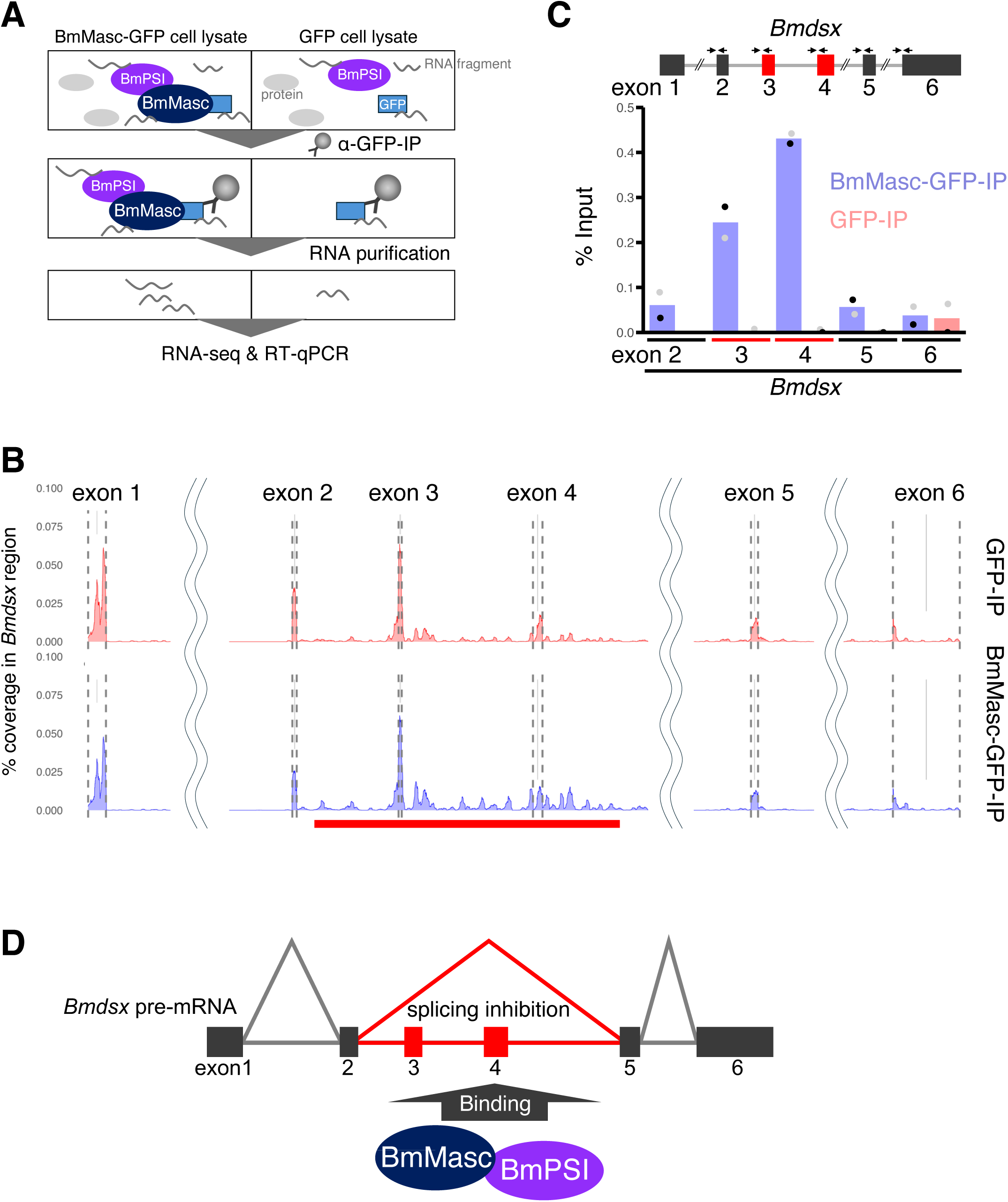
Physical association between BmMasc-containing protein complex and *Bmdsx* pre-mRNA. (**A**) Schematic representation of the RNA immunoprecipitation (RIP) experiments. The lysates of cells transfected with *BmMasc-GFP* or *GFP* (control) were immunoprecipitated with anti-GFP beads, and RNA fragments were purified from the immunoprecipitates and subjected to RNA-seq or RT-qPCR. (**B**) RIP-seq analysis of BmMasc-GFP-or GFP-associated RNA fragments. The x-axis indicates the genomic region of the *Bmdsx* gene, whereas the y-axis indicates the relative coverage in the *Bmdsx* region. The exon–intron boundaries are indicated by dashed lines. The region that was enriched by BmMasc-GFP-bound RNA fragments is highlighted by the red line. (**C**) RIP-qPCR analysis of BmMasc-GFP-or GFP-associated RNA fragments. The data are the means of two independent experiments. The positions of the primers used for RIP-qPCR are shown by arrows. (**D**) Proposed model of BmMasc-dependent *Bmdsx^M^* splicing. The BmMasc–BmPSI protein complex binds to the adjacent region of the female-specific exons of *Bmdsx* pre-mRNA and induces exon skipping, resulting in the production of *Bmdsx^M^*.

## Discussion

To fill in the missing piece in *B. mori* male determination process, we searched for BmMasc-interacting proteins and successfully identified BmPSI as the binding partner of BmMasc. Knockdown experiments clearly showed that BmMasc overexpression was insufficient to induce *Bmdsx^M^* expression under low-level expression of BmPSI, thereby demonstrating that BmPSI is essential for the BmMasc-dependent masculinization process. This finding aligns with a previous *in vivo* result, where *Bmdsx^M^* expression was suppressed in males of the *BmPSI* mutant strain generated using a binary transgenic CRISPR/Cas9 method (Xu et al., 2017). Collectively, these results strongly suggest that the BmMasc–BmPSI protein complex is the core machinery for the induction of *Bmdsx^M^* splicing.

A previous research using a mutant *B. mori* strain showed that *BmPSI* depletion resulted in a decrease in *BmMasc* mRNA, proposing a model in which *BmPSI* functions as the upstream regulator of *BmMasc* to increase its mRNA level (Xu et al., 2017). However, our experiments using *B. mori* cultured cells revealed that BmPSI physically interacted and cooperated with BmMasc in the masculinizing cascade. Further biochemical analyses using BmPSI derivatives demonstrated that AB motifs of BmPSI are essential for its interaction with BmMasc. Because AB motifs are conserved among PSI homologs in other lepidopteran insects (Wang et al., 2014; Wang et al., 2021), the interaction between BmMasc and BmPSI’s AB motifs is likely essential and common in the lepidopteran masculinizing pathway. We also discovered that deletion of one of the two CCCH zinc finger domains of BmMasc did not alter the association with BmPSI, suggesting that these domains are dispensable for BmPSI binding. This finding supports our previous results that the CCCH zinc finger domains of BmMasc are not required for the masculinization process in *B. mori* and BmN-4 cells (Katsuma et al., 2015; Kiuchi et al., 2019). The precise role of the BmMasc’s CCCH zinc finger domains remains unknown, warranting further investigation.

In *Drosophila*, the AB motif of PSI is required for its interaction with U1 snRNP-specific 70K (U1-70K) (Labourier et al., 2001). PSI inhibits the binding of U1 snRNP to the precise 5′ splice site through recruiting U1 snRNP to the pseudo-5′ splice site. According to the RIP results and the observation that the AB motifs of BmPSI are essential for its interaction with BmMasc, the following mechanism can be proposed: BmPSI inhibits female-type *Bmdsx* splicing through recruiting BmMasc (instead of U1-70K) to the regions around the junctions of intron 2 with exon 3 and intron 3 with exon 4 (i.e., CE1) of *Bmdsx*. Since exon 3 and 4 are specifically skipped in *B. mori* males (Ohbayashi et al., 2001; Suzuki et al., 2001), we concluded that the BmMasc–BmPSI complex binds to specific regions of *Bmdsx* pre-mRNA, blocking female-type splicing, and thereby promoting male-specific exon skipping. Higher-resolution analyses are needed to identify the precise regions of *Bmdsx* pre-mRNA that mediate the physical interaction with the masculinizing machinery containing BmMasc and BmPSI.

We previously identified the masculinizing domain of BmMasc, which contains two conserved cysteine residues, Cys-301 and Cys-304 (Katsuma et al., 2015). Mutations in these cysteine residues significantly reduced the BmMasc-dependent masculinizing activity in BmN-4 cells, suggesting that these residues are required for its structural conformation, stability, and/or interaction with cooperating partner proteins. In this study, we observed that mutations in the masculinizing domain did not affect the binding ability of BmMasc to BmPSI (Fig. 2G), indicating that the interaction between BmMasc and BmPSI alone is insufficient to induce *Bmdsx^M^* expression. We hypothesize that the BmMasc-centered masculinizing machinery may contain not only BmPSI but also additional proteins, some of which may bind to BmMasc via its masculinizing domain. Some of these unknown cofactors may be listed in our LC-MS/MS results. Future studies will focus on identifying the functional interactors of BmMasc involved in the masculinizing machinery, and the role of the masculinizing domain of BmMasc.

## Experimental procedures

### Cell culture, transfection, and quantitative RT-PCR (RT-qPCR)

The DNA fragments encoding *BmMasc-GFP*, *BmPSI-mCherry*, and their derivatives were cloned into the pIZ/V5-His-g3 vector (Hirota et al., 2021). BmN-4 cells were maintained at 26 °C in IPL-41 medium (Applichem, Germany) supplemented with 10% fetal bovine serum (FBS) (Gibco, USA). BmN-4 cells (4.0 × 10⁵ cells per 35-mm dish) were transfected with two types of plasmid DNAs (1 µg each) using FuGENE HD transfection reagent (Promega, USA). The culture medium was replaced with fresh medium at 24 h post-transfection. Total RNA was extracted from the transfected cells using TRI-REAGENT® (Molecular Research Center Inc., USA) at 3 days after transfection. cDNA was synthesized using avian myeloblastosis virus reverse transcriptase and an oligo-dT primer (TaKaRa, Japan), as described previously (Sugano et al., 2016). RT-qPCR for *Bmdsx^M^* and *ribosomal protein 49* (*rp49*) was performed using the KAPA™ SYBR FAST qPCR Kit (Kapa Biosystems Inc., USA), and their expression levels were calculated using the 2^-ΔΔCt^ method. The primers that were used for RT-qPCR are listed in Table S2.

### LC-MS/MS-based identification of BmMasc-interacting proteins

BmMasc-GFP, BmMascΔNLS-GFP, or GFP were transiently expressed in BmN-4 cells (4.0 × 10^6^ cells) seeded in 10 cm-diameter culture dishes. At three days post transfection, the cells were fixed with 0.1% formaldehyde, washed, and lysed on ice for 10 min in 1 mL of RIPA buffer [20 mM HEPES-KOH (pH 7.5), 1 mM EGTA, 1 mM MgCl_2_, 150 mM NaCl, 0.25% Na-deoxycholate, 0.05% SDS, and 1% NP-40], supplemented with a protease inhibitor cocktail cOmplete EDTA-free (Roche, Switzerland) and Benzonase (Merck, Germany). After centrifugation, the supernatants were incubated with GFP-Trap Magnetic Agarose (ChromoTek, Germany) for 3 h at 4°C under gentle rotation. The magnetic agarose beads were collected using a magnetic stand, washed four times with RIPA buffer, and then twice with 50 mM ammonium bicarbonate buffer. Proteins bound to the beads were digested by adding 200 ng of Trypsin/Lys-C mix (Promega) for 16 h at 37 °C. The digests were reduced, alkylated, acidified with trifluoroacetic acid (TFA), and desalted using a GL-Tip SDB (GL Sciences, Japan). The eluates were evaporated in a SpeedVac concentrator and dissolved in 0.1% TFA and 3% acetonitrile (ACN). The subsequent LC-MS/MS experiments were performed as described previously (Katsuma et al., 2022) using an EASY-nLC 1200 UHPLC connected to an Orbitrap Fusion mass spectrometer equipped with a nanoelectrospray ion source (Thermo Fisher Scientific).

The peptides were separated on a 75 μm inner diameter × 150 mm C18 reversed-phase column (Nikkyo Technos, Japan) with a linear 4–32% ACN gradient for 0–100 min, followed by an increase to 80% ACN for 10 min and a final hold at 80% ACN for 10 min. The mass spectrometer was operated in a data-dependent acquisition mode with a maximum duty cycle of 3 s. MS1 spectra were measured with a resolution of 120,000, an automatic gain control (AGC) target of 4e5, and a mass range from 375 to 1,500 m/z. Higher-energy collisional dissociation (HCD) MS/MS spectra were acquired in the linear ion trap with an AGC target of 1e4, an isolation window of 1.6 m/z, a maximum injection time of 35 ms, and a normalized collision energy of 30. Dynamic exclusion was set to 20 s. Raw data were directly analyzed against *B. mori* protein data (Kawamoto et al., 2022) supplemented with BmMasc-GFP and BmMascΔNLS-GFP sequences using Proteome Discoverer version 2.5 (Thermo Fisher Scientific, USA) with Sequest HT search engine. The search parameters were as follows: (a) trypsin as an enzyme with up to two missed cleavages; (b) precursor mass tolerance of 10 ppm; (c) fragment mass tolerance of 0.6 Da; (d) carbamidomethylation of cysteine as a fixed modification; and (e) acetylation of protein N-terminus and oxidation of methionine as variable modifications. Peptides and proteins were filtered at a false discovery rate (FDR) of 1% using the percolator node and the protein FDR validator node, respectively. Label-free precursor ion quantification was performed using the precursor ions quantifier node, and normalization was performed such that the total sum of abundance values for each sample over all peptides was the same.

### Immunoprecipitation and Western blotting

At three days post transfection, the BmN-4 cells were collected by scraping with disposable scrapers and lysed on ice for 15 min in 1 mL of chilled TNE-N buffer [20 mM Tris (pH 8.0), 1 mM EDTA, 150 mM NaCl, 1% NP-40] supplemented with cOmplete EDTA-free (Roche, Switzerland). After centrifugation at 20,000 × *g* at 4°C for 15 min, a fraction of supernatants was mixed with 2× SDS sample buffer [4% SDS, 20% glycerol (liquid), 125 mM Tris-HCl (pH6.8), 0.04% BPB, 10% 2-Mercaptoethanol] and boiled. The remaining supernatants were incubated with GFP-Trap Magnetic Agarose (ChromoTek, Germany) or RFP-Trap Magnetic Agarose (ChromoTek, Germany) for 1 h at 4°C under gentle rotation. The magnetic agarose beads were collected using a magnetic stand, washed three times with TNE-N buffer, and then mixed with 2× SDS sample buffer [4% SDS, 20% glycerol (liquid), 125 mM Tris-HCl (pH6.8), 0.04% BPB, 10% 2-Mercaptoethanol] and boiled to elute binding proteins. The lysate proteins and Eluted proteins were separated on 4–12% Bis-Tris gels (NuPAGE, Invitrogen, USA) in MOPS-SDS buffer (NuPAGE, Invitrogen, USA) using an XCell SureLock mini-cell (Invitrogen, USA). The proteins were transferred onto PVDF membranes in Transfer Buffer (NuPAGE, Invitrogen, USA) using an XCell II blot module (Invitrogen, USA), according to the manufacturer’s protocol. The membranes were blocked with 4% Block Ace (DS Pharma Biomedical, Japan), followed by incubation with the following primary antibodies: anti-GFP antibody (598, MBL, Japan), anti-BmPSI antibody (Suzuki et al., 2010), anti-RFP antibody (M204-3, MBL, Japan), or anti-Masc antibody (Kiuchi et al., 2019), in the antibody dilution buffer Kiwami Setsuyaku-kun (DRC, Japan). After incubation with the primary antibody, the membrane was washed four times with TBS-T buffer and incubated with the secondary antibody: HRP-bound anti-Rabbit IgG (111-035-144, Jackson ImmunoResearch Laboratories Inc., USA), or HRP-bound anti-Mouse IgG (626520, Invitrogen, USA). After incubation with the secondary antibody, the membrane was washed five times with TBS-T buffer and stained using a Immobilon Western Chemiluminescent HRP Substrate (Millipore, USA). Antibody-stained proteins were detected using a ChemiDoc XRS Plus imaging system (Bio-Rad, USA).

### Knockdown of *BmPSI* in BmN-4 cells

DNA fragments for ds*BmPSI* and ds*Luc* were amplified from *BmPSI-mCherry* and Bm31Luc (Nakanishi et al., 2010), respectively, using the primers listed in Table S2. Each dsRNA was transcribed from the DNA template using T7 Mega Script Kit (Thermo Fisher Scientific, Waltham, Massachusetts, USA). For dsRNA soaking experiments, BmN-4 sid-1 cells (Mon et al., 2012) were cultured at 26°C in TC-100 (Sigma-Aldrich, Missouri, USA) supplemented with 10% FBS (Gibco, USA) and tryptose phosphate broth (Sigma-Aldrich). BmN-4 sid-1 cells (1.0 × 10^5^ cells per 35-mm diameter dish) were soaked with three types of dsRNAs (4 µg each, twice). A second soaking was carried out three days after the first soaking. Three days after the second soaking, the cells are collected, lysed, and mixed with 2× SDS sample buffer, and then used for Western blotting to evaluate the knockdown efficiency. We used an anti-BmPSI antibody (Suzuki et al., 2008) and an anti-actin antibody (sc-1616-R, Santa Cruz Biotechnology) as primary antibodies, and HRP-bound anti-Rabbit IgG (111-035-144, Jackson ImmunoResearch Laboratories Inc., USA) as a secondary antibody. Three days after the second dsRNA soaking, the cells were transfected with 2 µg of *BmMasc-GFP* or *GFP* using FuGENE HD (Promega, USA), then collected and subjected to RT-qPCR three days after transfection.

### Identification of the interacting regions of BmMasc and BmPSI

Mutagenesis experiments were conducted using the KOD Plus Mutagenesis Kit (TOYOBO, Japan) according to the manufacturer’s protocol. The immunoprecipitation and Western blotting experiments were performed as described above.

### RNA immunoprecipitation (RIP) experiments

At three days post transfection, BmN-4 cells were collected by scraping with disposable scrapers and lysed on ice for 15 min in 1 mL of chilled TNE-N buffer (20 mM Tris pH 8.0, 1 mM EDTA, 150 mM NaCl, 1% NP-40) supplemented with cOmplete EDTA-free (Roche, Switzerland) and 200 U/µL of SUPERase (Thermo Fisher Scientific, USA). After centrifugation at 20,000 × *g* at 4°C for 15 min, 20 µL of the supernatants were mixed with TRI-Reagent. The remaining supernatants were incubated with GFP-Trap Magnetic Agarose (ChromoTek, Germany) for 1 h at 4°C under gentle rotation. The magnetic agarose beads were collected using a magnetic stand, washed three times with TNE-N buffer, and then mixed with 500 µL of TRI-REAGENT. Total RNA was purified according to the manufacturer’s protocol. We used 1 µL of glycogen (Roche Diagnostics, Switzerland) per tube as a coprecipitant for the RNA pellets. cDNA was synthesized using avian myeloblastosis virus reverse transcriptase with a random 9-mer (TaKaRa, Japan). RT-qPCR was performed using a KAPA™ SYBR FAST qPCR Kit (Kapa Biosystems Inc., USA), and % input values were calculated.

BmN-4 cells transfected with *BmMasc* or *GFP* cDNA were immunoprecipitated with anti-GFP beads using μMACS GFP Tagged Protein Isolation Kit (Miltenyi Biotec), as previously reported (Kiuchi et al., 2019). Immunoprecipitated RNA fragments were prepared from the beads and subjected to RNA-seq experiments. RNA-seq libraries were performed using Agilent Strand Specific library prep kit without poly(A) selection and analyzed on an Illumina HiSeq 2500 platform based on the manufacturer’s protocol for 100-bp paired-end reads. Raw RIP-seq reads were mapped to the *B. mori* genome (Kawamoto et al., 2019) using hisat2 (Kim et al., 2019). Subsequently, using samtools (Li et al., 2009), only the reads that mapped to the *Bmdsx* genomic region (from 10580681 to 10778244 in Bomo_Chr25) were extracted and converted to bed files using bedtools (Quinlan and Hall, 2010). Reads that mapped in the sense direction to the *Bmdsx* gene were then extracted, and the coverage of each base was calculated using coverageBed from bedtools. The libraries were normalized based on the total coverage of this region, and a graph was generated using the ggplot2, tidyr, and dplyr libraries in R (Wickham 2016; Wickham et al., 2024; Wickham et al., 2023).

## Data availability

All data presented in the figures of this paper are available upon on request. The RIP-seq data have been deposited in the DDBJ (DNA Data Bank of Japan) under accession numbers DRR318257–DRR318262. The MS proteomics data have been deposited to the ProteomeXchange Consortium via the jPOST partner repository with the dataset identifier PXD060754.

## Supporting information

This article contains supporting information.

## Supporting information

STable 1

STable 2

## Abbreviations

Bmdsx: Bombyx mori doublesex
BmIMP: Bombyx mori IGF-II mRNA binding protein
BmMasc: Bombyx mori Masculinizer
BmPSI: Bombyx mori P-element somatic inhibitor
dsRNA: double-stranded RNA
Fem: Feminizer
NLS: nuclear localization signal
piRNA: PIWI-interacting RNA
RIP: RNA immunoprecipitation
rp49: ribosomal protein 49

## Acknowledgments

We thank M. Kawamoto for his contribution of the initial stage of the project, K. Nishino for his help with LC-MS/MS analysis, and R. Hamajima and T. Kusakabe for providing BmN-4 sid-1 cells.

## Author contributions

T.Ka., and S.K. writing–original draft; Ta.K., N.M.-I., H.K., K.S., and S.K. investigation; T.Ki., and S.K. conceptualization; N.M.-I., H.K., K.S., and S.K. methodology; Ta.K., N.M.-I., H.K., K.S., and S.K. formal analysis; Ta.K., N.M.-I., H.K., K.S., M.G.S., Y.S., T.Ki., and S.K. writing–review and editing; M.G.S. resources; S.K. project administration; Y.S., T.Ki., and S.K. funding acquisition.

## Funding information

This work was supported by Grants-in-Aid for Scientific Research on Innovative Areas “Spectrum of the Sex: a continuity of phenotypes between female and male” (17H06431) to S. K. and T. Ki., Grant-in-Aid for Scientific Research (A) (22H00366) to S. K., G-7 Scholarship Foundation to S. K., and 16H06279 (PAGS) to Y. S.

## Conflicts of interest

The authors declare that they have no conflicts of interest with the contents of this article.

---

**Table S1. Proteins related to RNA splicing, as indicated in Fig. 1A**.

**Table S2. Primers used in this study.**

## Notes

### Competing Interest Statement

The authors have declared no competing interest.

