## Supplementary material for "The Masc–PSI complex directly induces male-type *doublesex* splicing in silkworms": STable 1

**Table S1. Proteins related to RNA splicing highlighted in Fig. 1A.**

| Accession | Abundance Ratio: BmMascGFP / GFP | Abundance Ratio: BmMascΔNLSmutGFP / GFP | Annotation |
|---|---|---|---|
| P_KWMTBOMO07543 | 0.368 | 0.249 | PREDICTED:_U4/U6.U5_tri-snRNP-associated_protein_1_isoform_X2_[Bombyx_mori] |
| P_KWMTBOMO06766 | 2.667 | 1.258 | PREDICTED:_splicing_factor_3B_subunit_3_isoform_X1_[Amyelois_transitella] |
| P_KWMTBOMO10443 | 0.237 | 0.233 | putative_alternative_splicing_type_3_and_[Danaus_plexippus] |
| P_KWMTBOMO10348 | 0.225 | 0.189 | putative_splicing_factor_pTSR1_[Danaus_plexippus] |
| P_KWMTBOMO02170 | 6.532 | 1.929 | PREDICTED:_pre-mRNA-splicing_factor_SYF1_[Amyelois_transitella] |
| P_KWMTBOMO07384 | 2.796 | 1.119 | putative_U2_snrnp_auxiliary_factor,_small_subunit_[Danaus_plexippus] |
| P_KWMTBOMO04690 | 8.745 | 1.765 | PREDICTED:_pre-mRNA-processing-splicing_factor_8_[Bombyx_mori] |
| P_KWMTBOMO15176 | 10.854 | 1.752 | PREDICTED:_pre-mRNA-processing_factor_39_isoform_X1_[Amyelois_transitella] |
| P_KWMTBOMO16286 | 5.014 | 0.659 | PREDICTED:_pre-mRNA-splicing_factor_38A_[Papilio_machaon] |
| P_KWMTBOMO06038 | 5.465 | 0.899 | PREDICTED:_pre-mRNA-processing_factor_19_[Bombyx_mori] |
| P_KWMTBOMO07485 | 0.358 | 0.288 | PREDICTED:_repressor_splicing_factor_1_isoform_X1_[Bombyx_mori] |
| P_KWMTBOMO14687 | 0.828 | 0.252 | PREDICTED:_putative_pre-mRNA-splicing_factor_ATP-dependent_RNA_helicase_PRP1_[Papilio_polytes] |
| P_KWMTBOMO01400 | 0.568 | 1.341 | Pre-mRNA-processing_factor_6_[Papilio_xuthus] |
| P_KWMTBOMO15210 | 5.76 | 1.694 | PREDICTED:_intron-binding_protein_aquarius_[Amyelois_transitella] |
| P_KWMTBOMO06801 | 0.738 | 0.394 | small_nuclear_ribonucleoprotein_protein_F_[Bombyx_mori] |

BmMasc-interacting proteins related to the following GOs are listed.

| | |
|---|---|
| GO:0008380 | RNA splicing |
| GO:0000398 | mRNA splicing, via spliceosome |
| GO:0005681 | spliceosomal complex |
| GO:0000245 | spliceosomal complex assembly |
| GO:0000387 | spliceosomal snRNP assembly |
| GO:0006376 | mRNA splice site selection |
| GO:0005685 | U1 snRNP |
