## Supplementary material for "The Masc–PSI complex directly induces male-type *doublesex* splicing in silkworms": STable 2

**Table S2. Primers used in this study.**

| Primer name | Sequence | Purpose |
|---|---|---|
| BmdsxM-F | CAAGGAAAATCTACGAAGGTTA | RT-qPCR |
| BmdsxM-R | GGTCATGCGCCGTCTGTATC | |
| Bmrp49-F | CCCAACATTGGTTACGGTTC | |
| Bmrp49-R | GCTCTTTCCACGATCAGCTT | |
| BmPSI-F | TACCGAGCTCGGATCATGAGTGATTATTCTTCTATGG | Cloning |
| BmPSI-R | GCCACTGTGCTGGATTCACTGCTGGTGGTC | |
| BmPSI-mCherry-R | GCTACCTCCTTGGATCTGCTGGTGGTCGG | |
| Bmdsx-2F | CTTCTTCCCTTTACGCATGCATAC | RIP-qPCR |
| Bmdsx-2R | GTGACAGTTCTCCACAAGCGTC | |
| Bmdsx-3F | AACGCAAAGGAACCTTTGCC | |
| Bmdsx-3R | TGCTTCCTCGCGTACTCGTC | |
| Bmdsx-4F | TCTATTCCACTAACAATGTACACTACA | |
| Bmdsx-4R | GACGAAGACAGTACACCAC | |
| Bmdsx-5F | CTGGCCTAAACGTATCACCGAC | |
| Bmdsx-5R | CGCCATTGATGCATCATCCAATAAC | |
| Bmdsx-6F | GGCGAGTACGGTTAGTTCGTTC | |
| Bmdsx-6R | GAAGTTGACGGGGAATCGGTG | |
| N-BmPSI-F | ATCCAAGGAGGTAGCGGTGGAG | Mutagenesis |
| N-BmPSI-R | AAAGTTGACAGGCCCACCAAC | |
| C-BmPSI-F | ATCCCGGAAGGCAACGGTAAC | |
| C-BmPSI-R | CATGATCCGAGCTCGGTACCAAG | |
| dAB1-C-BmPSI-F | GCTAAAATGCAGCAGCAGCAG | |
| dAB1-C-BmPSI-R | TATTGATACTTGTTGCTGCGGTCCCTG | |
| dAB2-C-BmPSI-F | ATTAAGCAGCAGCAAAACCAACAGCCGCAG | |
| dAB2-C-BmPSI-R | CGGGGTGGAGGGACCGC | |
| dzf1-BmMasc-F | CTAAAAGTTTTGAAAGAAACCGTAAAGTTTTGCCGCGATTTTC | |
| dzf1-BmMasc-R | TTTTTGTTCCGATGATGACGCGGTTATC | |
| dzf2-BmMasc-F | GAGGAGCAAAAGCTATTTG | |
| dzf2-BmMasc-R | CAAAACTTTTAGGTCTAGTTTGTGAAGGTG | |
| CS-BmMasc-F | AATAACAGTGTGCAGAGGGAACTCAGGATTG | |
| CS-BmMasc-R | GCTGTCGCTTTCTAAAGAATTCGCGTC | |
| dsBmPSI-1-F | TAATACGACTCACTATAGGGAATAGTCAAACGGCGGGATACG | in vitro transcription |
| dsBmPSI-1-R | TAATACGACTCACTATAGGGTGTGGAGCGTTCTCCCTTC | |
| dsBmPSI-2-F | TAATACGACTCACTATAGGGGCAGGGGAAACCCCATCAG | |
| dsBmPSI-2-R | TAATACGACTCACTATAGGGGTGCCAGTGCGGGTTGTAC | |
| dsLuc-F | TAATACGACTCACTATAGGGTTCCATCTGCCAGGTATCAG | |
| dsLuc-R | TAATACGACTCACTATAGGGCGTCCACAAACACAACTCCTC | |
